# SilencioMA: A single compound that targets both *MAPT* and *APP* as a potential disease modifying therapeutic for Alzheimer’s Disease

**DOI:** 10.64898/2026.11.09.765491

**Authors:** Paraskevi Sgourdou, Nicholas Vrettos, Kevt’her Hoxha, Thallyah Baffic, Edward B. Lee, Hong Xu, Virginia M-Y Lee, Zissimos Mourelatos

**Author notes:** Corresponding authors: Zissimos Mourelatos and Paraskevi Sgourdou.

## Abstract

Alzheimer’s Disease (AD) is the most common age-related neurodegenerative disease and the leading cause of dementia worldwide. AD pathogenesis involves aggregation of amyloid-beta (Aβ), produced from amyloidogenic proteolytic processing of Amyloid Precursor Protein (APP), and accumulation of misfolded, pathological tau (Microtubule Associated Protein Tau –MAPT–). Current efforts focus on developing disease modifying therapeutics (DMTs) against single targets: Aβ or tau. However, DMTs that concurrently target both tau and Aβ are likely to have the greatest impact since they target both culprits of AD pathogenesis. Here we describe the development of SilencioMA, a single compound that targets both *MAPT* and *APP* messenger RNAs (mRNAs) based on <u>silen</u>cing <u>c</u>onnected interfering <u>o</u>ligonucleotides against *<u>M</u>APT* and *<u>A</u>PP*. Cerebrospinal fluid (CSF) administration of SilencioMA reduces Aβ and tau and pathology after a single intracerebroventricular injection in 5XFAD/PS19 mice, an animal model of severe amyloidosis and tauopathy. SilencioMA does not rely on lipid nanoparticles or lipophilic conjugates for CSF administration. At the same time its modular design permits conjugation of ligands that facilitate transcytosis through the blood brain barrier, allowing systemic administration. Our findings provide proof-of-concept for a strategy that targets the dual Aβ and tau pathologies of AD, and establishes a compelling foundation for further development of SilencioMA as a next-generation DMT for AD.

## INTRODUCTION

Alzheimer’s Disease (AD) is the leading cause of dementia afflicting an estimated 7.2 million people in the US aged 65 and older, which is expected to increase to 13.8 million by 2060 [1]. The causal substrate of AD is the underlying neuropathologic change (ADNC), which develops while people are asymptomatic, with clinical symptoms appearing as ADNC burden increases [45]. ADNC is detected with imaging and fluid biomarkers, which are effective for presymptomatic diagnosis and for biological and clinical staging as AD progresses [38] [45]. ADNC consists of extracellular amyloid plaques, and intracellular neurofibrillary tangles (NFTs), pre-tangles and neuropil threads (NTs) [8] [43] [67] [21]. Amyloid plaques consist of fibrillar aggregates of amyloid-beta (Aβ) peptides, derived from amyloidogenic proteolytic processing of Amyloid Precursor Protein (APP) [34] [61] [49] [37]. Aβ fibrils may aggregate in blood vessels, giving rise to Cerebral Amyloid Angiopathy (CAA), which often accompanies AD [14] [21]. NFTs and NTs consist of hyperphosphorylated tau fibrils (Microtubule Associated Protein Tau –MAPT) [93] [36] [54] [57] [87] [18]. A subset of amyloid plaques is intimately associated with dystrophic and swollen neurites containing phosphorylated tau fibrils and are known as neuritic plaques (NPs) [43] [67] [21]. The pathogenesis of AD, and of most age-related neurodegenerative diseases, involves protein misfolding into toxic conformers known as amyloid fibrils/ filaments [22] [78] [33] [18]. Aβ and tau fibrils are the molecular substrates of AD, while tau fibrils alone underlie tauopathies [29] [17] [94] [78] [18].

Therapeutic strategies that target tau and Aβ are expected to be disease-modifying since they target the proximal causes of AD pathogenesis [47] [16] [12] [55] [45] [80] [75] [41]. They can be broadly categorized as strategies that promote clearance of pathological proteins or strategies that reduce their production. Passive immunotherapies represent the largest class of AD therapeutics that target the pathological proteins for clearance [47] [16] [12] [55] [80] [41]. Two Food and Drug Administration (FDA)-approved, anti-Aβ monoclonal antibodies (mAbs) (Lecanemab and Donanemab) efficiently remove amyloid plaques, slowing cognitive decline by up to ∼27% in patients with early AD [80]. However, anti-Aβ therapy may provoke adverse events, known as Amyloid Related Imaging Abnormalities (ARIA) associated with brain edema (ARIA-E) or brain microhemorrhages (ARIA-H); these adverse events are more common and more severe in patients with CAA and/or carrying apolipoprotein E4 (*APOE4*) alleles [47] [12] [30] [52]. Moreover, anti-Aβ therapies do not target pathological tau, which is the main driver of neurodegeneration, and it is unlikely that they will be efficacious in patients that have advanced past mild dementia [47] [16] [5]. Targeting tau with mAbs is more challenging than targeting Aβ, as tau is predominantly intracellular. The first generation of anti-tau mAbs failed in clinical trials of AD and tauopathies; second generation mAbs are in clinical trials either alone or in combination with anti-Aβ mAbs [16] [12].

Another promising strategy is to reduce production of Aβ and tau proteins by targeting their mRNAs (*APP* and *MAPT*, respectively) using antisense oligonucleotides (ASOs) or small interfering RNAs (siRNAs) [86] [25] [53] [11] [19] [23]. This strategy is effective for both extracellular and intracellular targets (like tau). An ASO against *MAPT* (BIIB080) [65], administered in CSF via intrathecal injection for early AD, showed reduction in tau pathology and cognition benefit in the Phase 2 CELIA trial, although the biomarker response was essentially flat across the dose range and did not meet the study’s primary endpoint, which was to demonstrate a dose–response relationship (NCT05399888).

siRNAs, by entering the endogenous microRNA pathway and assembling with the Ago2 nuclease, form multiple turnover enzymes, which are more specific and potent than ASOs [42] [59] [64]. Chemical modifications, including substitution of all 2’ hydroxyl groups of ribose sugars, replacement of phosphodiester bonds with phosphorothioates at select positions, and use of conjugates, has led to their development as powerful pharmacological agents [2] [15] [48] [79] [3] [13] [23] [84]. siRNAs targeting *MAPT* are in early phases of clinical trials with LY3954068 (Eli Lilly) delivered intrathecally; and ARO-MAPT (Arrowhead Pharmaceuticals) and DNL-628 (Denali) administered systemically for AD and tauopathies as siRNA-brain shuttle conjugates [7] [39]. Notably, targeting *MAPT* with siRNAs led to dose-dependent reduction of tau in CSF without adverse effects in healthy volunteers [4]. An siRNA targeting *APP* (mivelsiran, Alnylam), delivered intrathecally, is in phase 1 trial for AD and CAA [6].

Aβ and tau pathologies are closely associated, suggesting that single-target approaches, whether directed at APP/Aβ or MAPT/tau in isolation, may not be sufficient for disease modification across the full spectrum of AD. A therapeutic approach capable of simultaneously suppressing both molecular drivers of neurodegeneration may modify AD in a way that monotherapy cannot. To date, there are no AD therapeutics capable of reducing both Aβ and phosphorylated tau simultaneously. Here, we describe the design, synthesis, and evaluation of a novel, dual-targeting siRNA therapeutic that we term <u>Silen</u>cing <u>c</u>onnected interfering <u>o</u>ligonucleotide against *<u>M</u>APT* and *<u>A</u>PP* (SilencioMA). SilencioMA simultaneously silences both *APP* and *MAPT*, leading to reduction of Aβ and phosphorylated tau as well as amelioration of pathology in a triple transgenic AD mouse model that develops severe ADNC. By co-targeting the two core drivers of AD pathology at the mRNA level, SilencioMA aims to overcome limitations of current monotherapies, while preserving the favorable tolerability profile that distinguishes siRNA approaches from immunotherapies. Our findings provide proof-of-concept for a strategy that targets the dual Aβ and tau pathologies of AD and establishes a compelling foundation for further development of SilencioMA as a next-generation DMT for AD.

## MATERIALS AND METHODS

### Mouse lines

All mouse experiments were approved by the Institutional Animal Care and Use Committee (IACUC) of the University of Pennsylvania and were performed in accordance with the ethical guidelines of the National Institutes of Health (NIH). Mice were raised under a constant 12-hour light/dark cycle. The mice used in this study were produced by crossing PS19 (B6.Cg-Tg(Prnp-MAPT*P301S)PS19Vle/J, Strain: 024841), with 5XFAD (B6SJL-Tg(APPSwFlLon,PSEN1*M146L*L286V) 6799Vas/Mmjax), both lines purchased from Jackson Labs. All offsprings were genotyped for the presence of transgenic *MAPT, APP, PS1*. Both male and female mice, positive for all three transgenes, were used in this study.

### Oligonucleotides used for gene silencing

For the initial *in cellulo* screens, siRNAs were synthesized by Integrated DNA Technologies (IDT). Preclinical/therapeutic grade modified oligonucleotides for SilencioMA were synthesized and purified by Eurogentec. Lyophilized oligonucleotides were resuspended in artificial CSF (aCSF) buffer (137 mM NaCl, 5mM KCl, 20mM D-(+)-Glucose, 8mM HEPES, pH 7.4). To generate SilencioMA, one part of M2024-Guide and one part of A2185-Guide were annealed to one part of MA-passenger by heating at 65° C for 5 minutes, followed by 37° C for 60 minutes and cooled on bench to room temperature. Stock concentration of SilencioMA was at 8mM; dilutions were made with aCSF.

### Stereotactic intracerebroventricular (ICV) injections

The mice underwent bilateral, intracerebroventricular (ICV) injection by insertion of a single needle into the brain at established coordinates from Bregma using a stereotactic rack: +0.2mm to +0.3 mm Anterior, −1.00 mm Lateral (right) and +1.00mm Lateral (Left), −2.5mm to −3.00mm Ventral. Each of the two injection sites received 5μl of 4mM SilencioMA, or aCSF only. The procedure was as described in [40]. Briefly, mice were deeply anesthetized with ketamine/xylazine/acepromazine, immobilized in a stereotaxic frame (David Kopf Instruments). The injection on either site was performed at a rate of 0.4 μl/min followed by a 2min wait before the needle was pulled out. The body temperature of the mice was maintained at 37 °C with a heating pad during surgery while subsequently they were placed under a heating lamp and monitored until awake and and exhibiting voluntary locomotion (≥ 2 hours). Mice were also monitored for a minimum of 3 days after surgery.

### Cell culture and transfection

HEK-293T cells and SHSY5Y cells were maintained in Dulbecco’s modified Eagle’s medium (DMEM) and DMEM/F12, respectively, at 37 °C and 5% CO_2_. Media were supplemented with 10% fetal bovine serum (FBS) (ThermoFisher Scientific, Waltham, MA, USA) and 1% Glutamine. Transfections were performed with RNAiMAX Transfection Reagent (ThermoFisher Scientific, Waltham, MA, USA) according to manufacturer’s instructions. The cells were harvested 48h after transfection.

### RNA/protein extraction

Transfected cells were processed with Qiagen RNeasy mini kit for RNA extraction and subsequent Reverse Transcription-quantitative Polymerase Chain Reaction (RT-qPCR) assays. Frozen brain tissue was ground with plastic pestle in RIPA buffer (50 mM Tris-HCl, pH 7.5, 150 mM NaCl, 5 mM EDTA, 1% NP-40, 0.5% sodium deoxycholate, 0.1% SDS) supplemented with RNAse (1u/ul), protease and phosphatase inhibitors (1X), then sonicated 3 times at 30% output. Half of the sonicated lysate was mixed with Trizol and used for RNA purification with miRNeasy kit (Qiagen). The remaining lysate was centrifuged at 16.000g for 15 min, at 4°C. The amount of protein in the samples was measured by Bradford assay, and lysates were stored at −80°C.

### mRNA quantification

250ng of total RNA were reverse transcribed using the Quantitect kit (Qiagen). The produced cDNAs were amplified with PowerUp SYBR-green mix (Applied Biosystems) and the primers used are listed in Supplementary Table 1. Assays were run on StepOnePlus Real-Time PCR system (Applied Biosystems) and quantified with the ΔΔCt method [60] *Gapdh* mRNA was used for input normalization. Results were calculated from 3 technical triplicates and analyzed using Graphpad-Prism 10.4.1

### Meso Scale Discovery (MSD) Electrochemiluminescent Immunoassays

The total amount of tau protein and the phosphorylated form pT231 present in each sample lysate were quantified using the MSD electrochemiluminescent immunoassay (Phospho(Thr231)/Total Tau Kit, catalog number K15121D). For the quantification of β-Amyloid species in each sample, the V-PLEX Plus Aβ Peptide Panel 1 (6E10) kit was used (catalog number K15200G-1). Samples were prepared according to the manufacturer’s protocol, involving centrifugation followed by dilution with kit-specific sample diluent to reach the Minimum Required Dilution (MRD). Each sample, along with the appropriate negative and positive controls, was run in technical duplicates and read on MESO QuickPlex SQ 120MM instrument.

### *In Situ* Hybridization (ISH) with Digoxigenin-labeled Locked Nucleic Acid (LNA) probes

The procedure was adopted from [68] with few modifications. Briefly, we designed LNA probes against M2024 (5’-AG+GTGGCCA+GGT+GGAAG+TAAA, +denotes LNA base) and A2185 (5’-TGCT+GAA+GAA+GA+AACA+GTACA), synthesized at IDT. The probes were digoxigenin labeled with Dig-ddUTP and the Dig-3’-end labeling kit (Roche) and stored at −20° C until use. 5 μm-thin sections of FFPE tissue were placed onto Probe-on slides. Slides were deparaffinized in xylenes, 2 x 5 min followed by 5 min each in serial dilution of ethanol (100%, 100%, 75%, 50%, 25%), followed by 2 changes of ddH_2_O. Unmasking of tissue RNA was done using Heat induced epitope (antigen) retrieval (HIER) using basic buffer (10 mM Tris-base, 1 mM EDTA solution, pH 9.0) from Vector Labs (H-3301-250). 1X Basic-Hier solution was pre-heated in a glass beaker on a hot plate until temperature reached 95-100 °C. The slides were immersed in staining rack in the heated buffer for 10min (up to 15 min) and then cooled down at RT. Subsequently, slides were digested with proteinase K solution (1X) for 10 min at 37° C, rinsed in PBS (2 rinses) and incubated with Hybridization mix (miRNA ISH buffer and LNA Detection Probes) for 1h at 55°C. Sections were subjected to stringent washes at 55°C in 5xSSC (1time), in 1xSSC (2 times), 0.2xSSC (2 times) and 0.2xSSC (1 time) at RT before transferring to PBS. Sections were blocked for 1hr in blocking solution (2% sheep serum, 2mg/ml BSA in PBST) at RT followed by incubation with a sheep anti-DIG-AP antibody (Roche catalog # 11 093 274 910, dilution 1:1000) at 4°C overnight in a humidified chamber. Next, slides were washed 4 times, 10 min each in PBST, and the AP reaction was achieved by incubating the slides with 1-Step NBT/BCIP plus Suppressor for approximately 2hours at 30°C in the humidifying chamber protected from light. KTBT buffer was used 2 times for 5 min each to stop the reaction. Slides were washed with water, 2 times for 1 min each, and counterstained with Nuclear Fast Red for 1min. After several changes of H_2_O, the sections were dehydrated by passing through a series of alcohols (50%, 75%, 95%, 100%, 100%) and xylenes and coverslipped in Cytoseal.

### Histology and Immunohistochemistry (IHC)

Adult mice were intracardially perfused with PBS, their brains were removed and mid-sagitally transected into two hemispheres. One hemisphere was immediately submerged in a solution of 10% Neutral Buffered Formalin (NBF) for fixation overnight at 4°C, and the second one was dissected in cold PBS, flash frozen in liquid nitrogen or dry ice for biochemical experiments. The fixed hemisphere was further sliced with a mouse brain slicer and processed for embedding in paraffin blocks. Sections were deparaffinized 2 x 5 min in xylene and immersed in descending EtOH series (2 x 100%, 1 x 95%, 1 x 80%, 1 x 70%) 1 min each. Antigen retrieval was done with Formic Acid (88%) for 5 min, then rinsing with water for 5 min. Slides are then immersed in freshly prepared H_2_O_2_/water (50 mL H_2_O_2_ to 150 mL water/boat) for 30 min followed by wash under running tap water for 10 min and in 0.1M Tris pH 7.6 for another 5 min. Blocking is done with 0.1 M Tris/2% FBS (PH=7.6 needs to be confirmed before use) for 5 min. followed by incubation in Ab solution AT8 (Phospho-Tau Ser202, Thr205, AT8 monoclonal, Thermo Fisher Scientific MN1020, 1:1000) or NAB228 (beta Amyloid (N-terminal) Monoclonal Antibody (NAB228) - 37-4200 Thermo Fisher Scientific, 1:1000) overnight at 4°C in a humidified chamber. After washing in 0.1M Tris pH 7.6 for 5 min, we block in 0.1 M Tris/2% FBS for another 5 min before incubating in secondary biotinylated anti-mouse antibody 1:1,000 in 0.1 M Tris/2% FBS for 1 hr. at RT. We repeat washing and blocking for 5min each, followed by incubation in AB solution for 1 h at room temperature. The sections are incubated in DAB for 8min or 4min, for AT8 or NAB228 Abs, respectively (time is kept consistent in all stainings), followed by washes in 0.1 M Tris pH 7.6 for 2 x 2 min and counterstained with Harris Hematoxylin solution for 20 seconds. After washing the sections in running tap water for 10 min we dehydrate in ascending series of ethanol (1 x 70%, 1 x 80%, 1 x 95%, 2 x 100%), 1 min each, and clear in 2 changes of xylene, 5 min each, before coverslipping in Cytoseal. The slides were scanned with 3DHistech Pannoramic 250 while AT8 and NAB228 expression was quantified with QuPath software, version 0.6.0 [9].

### Statistical analysis

Statistical analysis was performed using unpaired, one-tailed Welch’s t-test, or unpaired Mann-Whitney one-tailed test, depending on whether the data followed normal (Gaussian) distribution or not, respectively. A one-tailed test was specified *a priori* because SilencioMA is designed to lower mRNA levels of *MAPT* and *APP*, so only a reduction relative to vehicle (aCSF) was of interest. Analyses and graphing were performed with Graphpad-Prism 10.4.1. The Z-factors [97] calculated for *MAPT* mRNA and *APP* mRNA RT-qPCR assays are 0.85 and 0.90, respectively. Data in the graphs are shown as box plots (median, interquartile range, min–max whiskers; diamond = mean). For MSD immunoassays, one aCSF-treated animal was excluded from all MSD analyses (total tau, pT231-tau, Aβ38, Aβ40 and Aβ42) because its posterior cortex lysate showed total tau and pT231-tau levels 4.3-fold and 10.8-fold higher, respectively, than the mean of the remaining controls, and was identified as a significant outlier by Grubbs’ test (α = 0.05). Because all MSD assays were performed on the same lysate, the animal was excluded from the Aβ analyses as well. MSD analyses therefore include n = 14 for SilencioMA-treated and n = 7 for aCSF-treated mice. MSD values for all animals, including the outlier, are reported in Supplementary Table 3.

## RESULTS

### Design and generation of SilencioMA

To simultaneously target both *MAPT* and *APP* mRNAs we used siRNAs [25], which enter the endogenous miRNA pathway and assemble with the Ago2 nuclease to form multiple turnover enzymes that potently silence target genes [42, 59, 64] (Figure 1a). We first designed individual siRNAs against *MAPT* and *APP* using the following criteria: a) optimal design principles that promote loading of the guide strand to Ago2, target accessibility and specificity (reviewed in [89]; b) the ability to target and silence all human *MAPT* and *APP* mRNA isoforms; c) an ability to target *MAPT* and *APP* homologs of multiple species: human, mouse, rat, cynomolgus macaque; and d) the ability to target in addition to the wild-type human *MAPT* and *APP* alleles, all major disease-associated variants. We tested several siRNAs *in cellulo*, in human 293T cells, which have been shown to have neuronal progenitor-like properties [81] and in SH-SY5Y human neuroblastoma cells [10]. We chose M2024 (against *MAPT*) and A2185 (against *APP*) for further development as fully modified duplexes, which we will refer to as small interfering oligonucleotides (siOs), since all 2’-hydroxyl groups were substituted with either 2’-Fluoro or 2’-O-methyl groups. We covalently linked the passenger (sense) strands of the two siOs, in a 5’ to 3’ manner, by using polyethylene glycol linker, and annealed the two guide (antisense) strands, M2024 and A2185, to generate SilencioMA (Figure 1b). The guide strands contain 5΄-(E)-vinylphosphonate (5’-E-VP), rendering them resistant to phosphatases and promoting Ago2 binding [74] [26]. Except for 5’-E-VP, all chemical modifications that we employ (Figure 1c) are used in FDA-approved siRNAs, promoting stability and reducing immunogenecity [23]. We unpaired the first two nucleotides between the 5’ end of the guide strand and the passenger strand to decrease 5’-terminal thermodynamic stability, which is expected to facilitate Ago2 loading of the guide strand [91]. By connecting the passenger strands of SilencioMA, we enable equal uptake of the guides (that target *APP* and *MAPT*) by the same cells. Furthermore, linking the passenger strands renders them long and disfavor loading to Ago2, thereby reducing any off-target effects from the passenger strand. All sequences, with modifications, used in the production of SilencioMA are shown in Table 1. The siOs used to generate SilencioMA were purified by Ultra Performance Liquid Chromatography (UPLC) and their identities were confirmed by Mass Spectrometry (MS); representative chromatographs and mass spectra are shown in Supplementary Figure 1. To test the efficacy of SilencioMA in downregulating both MAPT and APP transcripts, we calculated its half maximal inhibitory concentrations (IC50) from 8-point dose-response curves using human 293T cells. We found that SilencioMA is highly potent with IC50s of 23pM for *MAPT* and 26pM for *APP* (Figure 1d).

**Figure 1.**
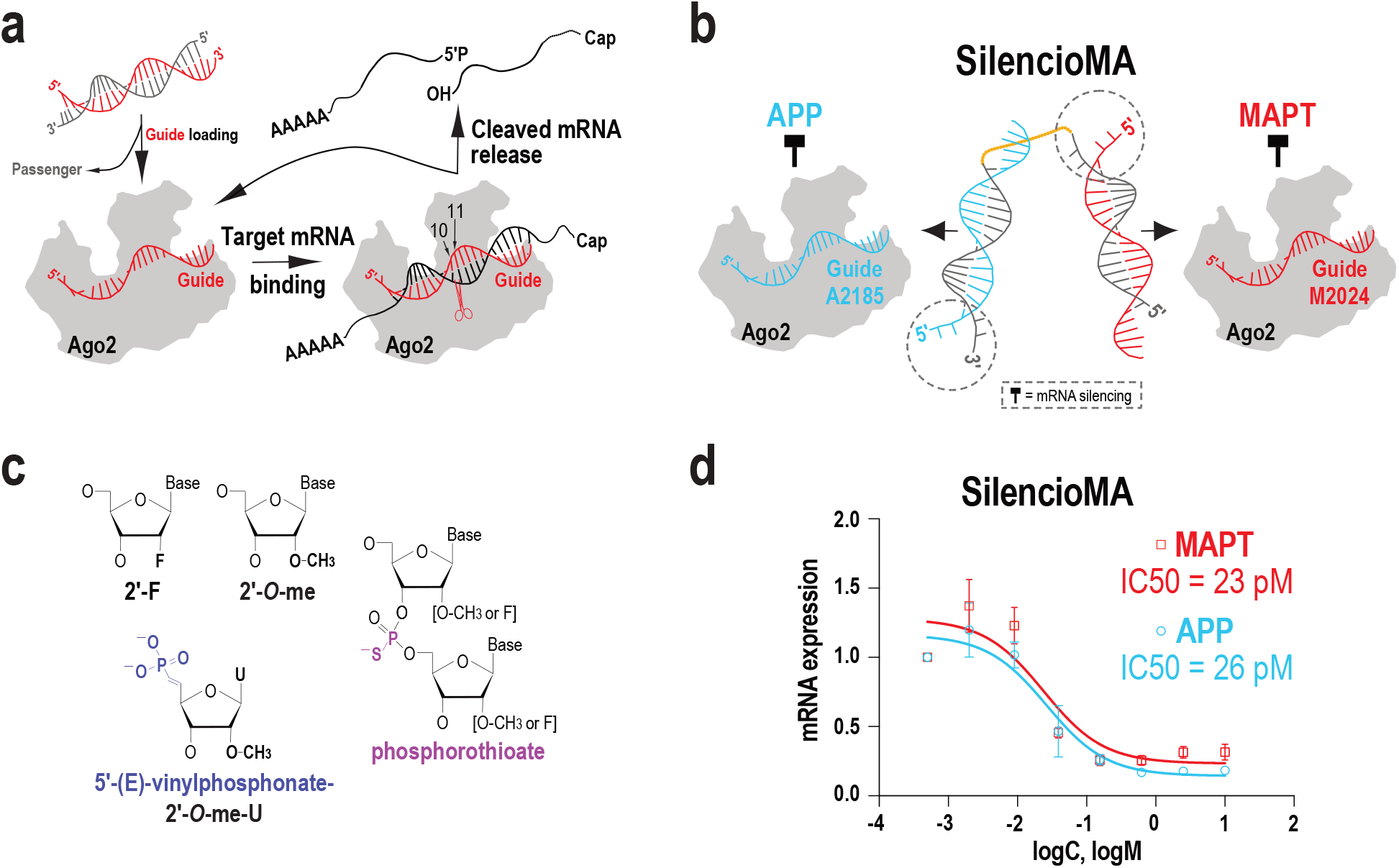
Design and properties of SilencioMA. **(a)** Schematic of RNA silencing by Ago2 and siRNAs. The guide (antisense) strand of an siRNA duplex is loaded to the Ago2 protein while the passenger (sense) strand is released. Guide RNA recognizes its target mRNA with antisense complementarity, triggering the Ago2 nuclease domain to cleave the phosphodiester bond of the target mRNA at a position across the 10th and 11th nucleotide of the guide RNA (scissors). The cleaved mRNA is released and further degraded by cellular exonucleases. The Ago2-Guide ribonucleoprotein is then available for further rounds of target mRNA recognition and cleavage, leading to potent gene silencing. **(b)** Schematic of SilencioMA: the linker that covalently connects the 5’ end of one passenger strand to the 3’ end of another passenger strand, is shown in orange. Dashed circles show the two unpaired bases between guide and passenger strand. The duplex is generated by annealing equal parts of M-passenger strand (grey), A2185-Guide (blue) and M2024-Guide (red). **(c)** Schematic of the modifications used. **(d)** Half-maximal inhibitory concentration (IC50) of SilencioMA, in human 293 cells, calculated from 8-point dose response curves performed in triplicates (10 nM, 2.5 nM, 0.625 nM, 0.156nM, 0.039 nM, 0.009 nM, 0.002 nM, 0.0006 nM).

**Table 1.** Sequences and modification of oligonucleotides used for SilencioMA.

|  |  |
|---|---|
| **M2024-Guide:** | vPmU*fU*mUfAfCfUmUfCmCfAmCfCmUfG*mG*fC*mC*mA*mC*mC*mU |
| **A2185-Guide:** | vP-mU*fG*mUfAfCfUmGfUmUfUmCfUmUfC*mU*fU*mC*mA*mG*mC*mA |
| **MA-Passenger:** | 5OH-mG*mC*mCfAmGmGmUfGmGmAmAmGmUfA*mU*mU-L36-mA*mA*mGfAmAmGmAfAmAmCmAmGmUfA*mG*mU |
| **SilencioMA:** | M2024-Guide + A2185-Guide + MA-Passenger, in 1:1:1 ratio |

(m = 2'-O-me;  f = 2'F; vPmU = Uridine with 5' E-vinylphosphonate and 2'-O-me; \*= phosphorothioate; L36 =polyethylene glycol linker containing 36 atoms)

### SilencioMA shows widespread distribution and retention in the mouse brain

To test the efficacy of SilencioMA in reducing tau and Aβ pathology, we crossed 5XFAD mice with PS19 mice to generate 5XFAD/PS19 triple transgenic mice that develop early ADNC [77]. PS19 express the T34 isoform (1N4R) of human *MAPT* with a P301S pathogenic variant; the mutant tau transgene is expressed at ∼5-fold higher levels than endogenous mouse tau, and PS19 are a widely used model for tau pathology [96]. The 5XFAD double transgenic mouse model (B6SJL-Tg(APPSwFlLon,PSEN1*M146L*L286V) expresses human *APP* with 4 mutations found in familial AD (FAD) patients (K670N, M671L, I716V, V717I) and human *PSEN1* with 2 FAD mutations (M146L and L286V). 5XFAD develops abundant Aβ pathology, with robust accumulation of amyloid plaques from 2 months onwards, and is among the most widely used model of amyloidosis [71] [24] [72] [31].

We ICV-injected a cohort of 5XFAD/PS19 mice (2 months old) with 40nmoles of SilencioMA (n=14) dissolved in artificial Cerebrospinal Fluid (aCSF), a buffer that mimics the electrolytes, osmolarity, and pH of native CSF. As controls, we injected mice with vehicle (aCSF) only (n=8). All mice were sacrificed eight weeks after injection, intracardially perfused with PBS and their brains dissected. One hemisphere was fixed in formalin and processed for histology, while the second was further dissected in cold PBS and flash frozen for biochemical studies (Figure 2a). To examine the cellular distribution of SilencioMA, we employed *in situ* hybridization (ISH) using Locked Nucleic Acid (LNA) modified probes that we designed against the guide strands (M2024 and A2185). We found widespread cellular distribution of both M2024 and A2185 guides across the entire 5XFAD/PS19 mouse brain (Figure 2b-g), confirming the retention of SilencioMA for at least 2 months post injection.

**Figure 2.**
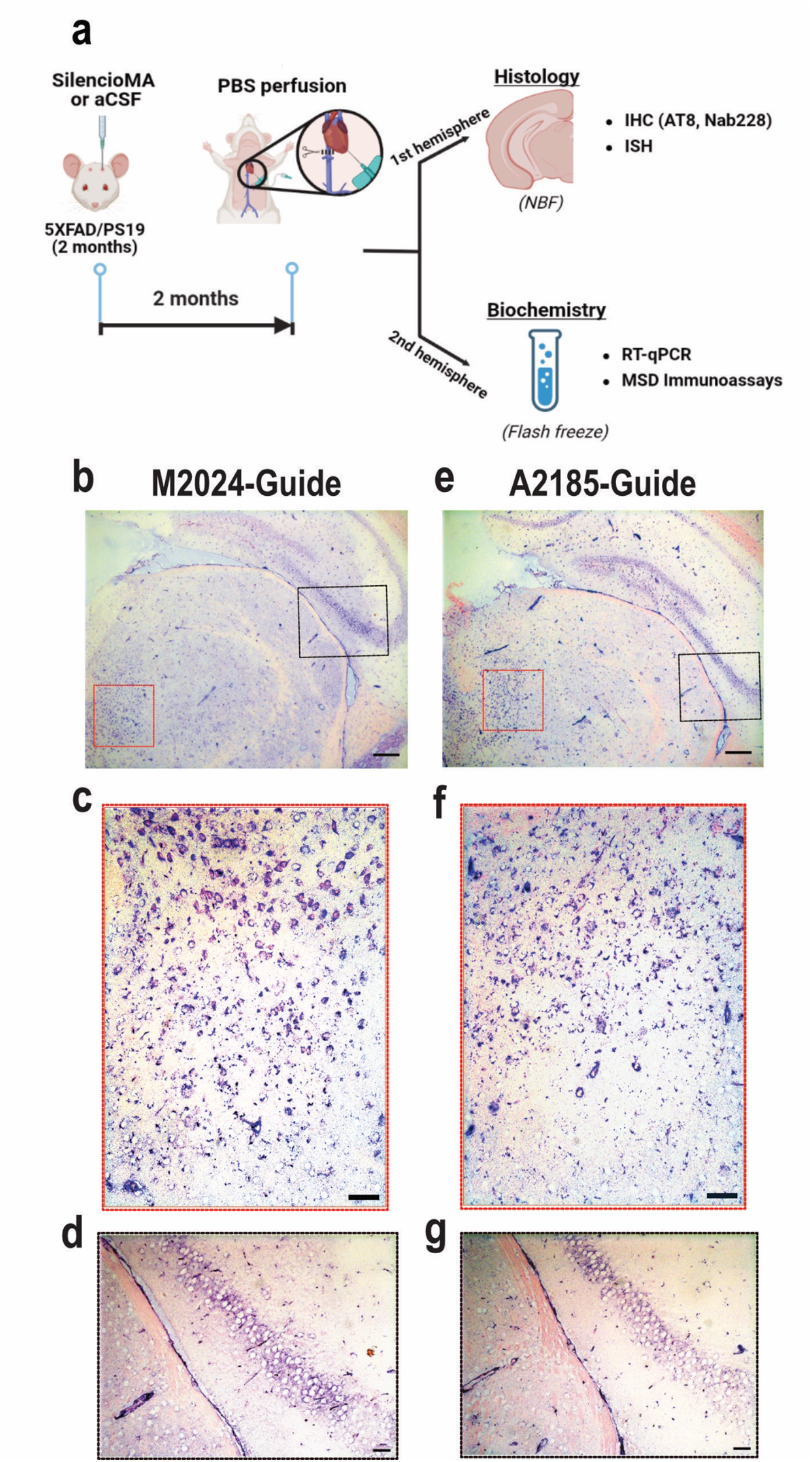
Widespread distribution and cell retention of SilencioMA. **(a)** Experimental workflow. Representative images of LNA-ISH against M2024 (**b, c, d**) and A2185 (**e, f, g**) in 5XFAD/PS19 mouse brains shows widespread cellular uptake and distribution of M2024 and A2185 (purple chromogens). Higher magnification images (**c, d, f, g**) correspond to boxed areas from the images shown above (**b** and **e**); nuclear red = counterstain; scale bars: b, e = 500μm, c, d, f, g = 50μm. Panel a was created with BioRender.com.

### SilencioMA significantly reduces *MAPT* and *APP* mRNAs, as well as tau, pT231-tau, and Aβ peptides in the 5XFAD/PS19 mouse model of AD

To assess functional target engagement by SilencioMA, we developed RT-qPCR assays for quantitation of human *MAPT* and *APP* mRNAs expressed from the transgenes (Supplementary Table 2). We found significant reduction of both human *MAPT* and *APP* mRNAs in the posterior cortex and hippocampus of SilencioMA injected mice compared to controls. *MAPT* mRNA levels were reduced by 26.1% (p=0.002) in posterior cortex and by 21.5% (p=0.037) in hippocampus (Figure 3a). *APP* mRNA levels were reduced by 25.4% (p=0.009) in posterior cortex and by 27.7% (p=0.010) in hippocampus (Figure 3b).

**Figure 3.**
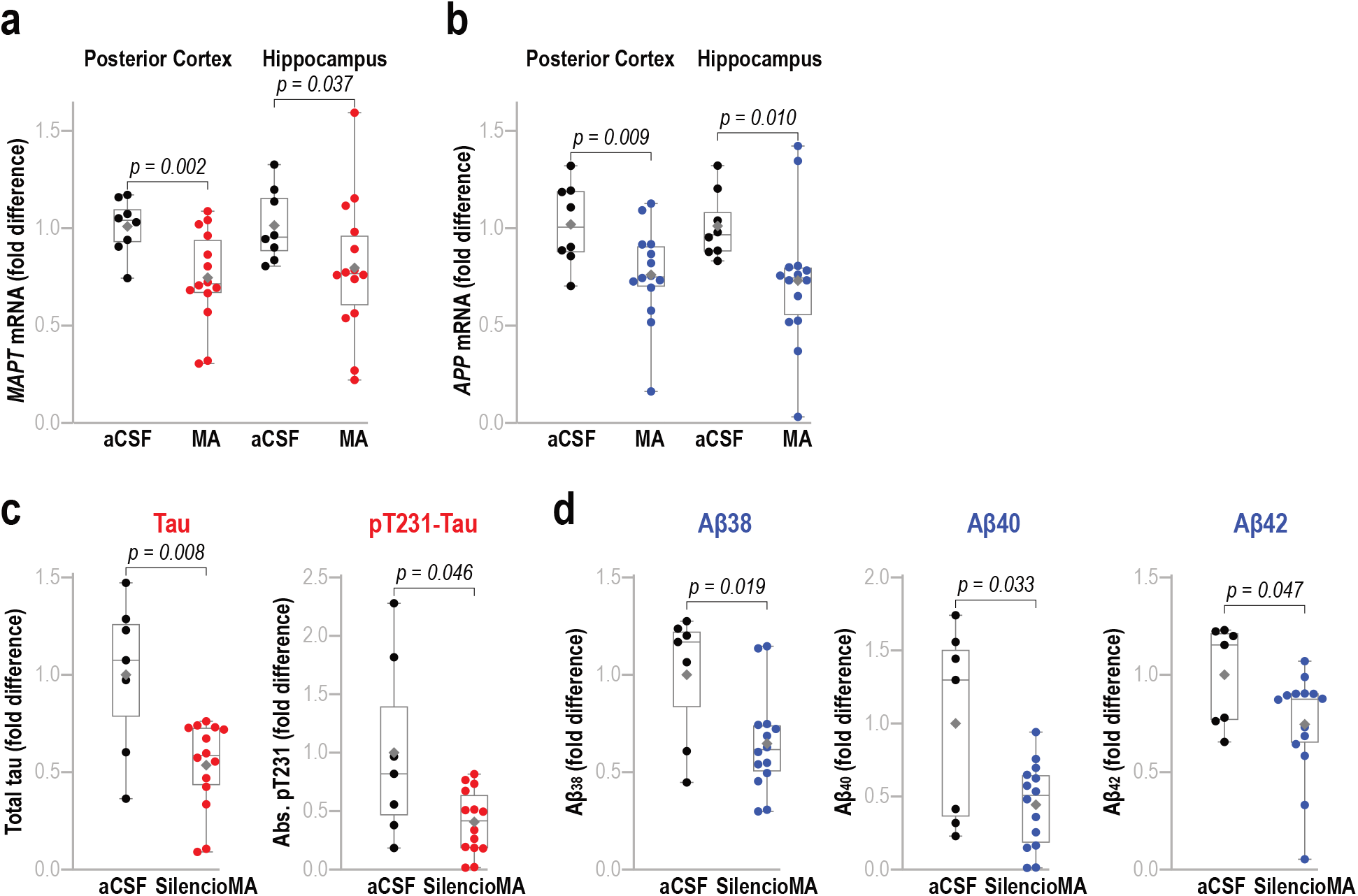
SilencioMA significantly reduces *MAPT* and *APP* mRNAs, tau, pT231-tau, and Aβ peptides in 5XFAD/PS19. Box plots of the fold difference of human *MAPT* (**a**) and human *APP* (**b**) mRNAs in SilencioMA compared to aCSF injected 5XFAD/PS19 mice in indicated brain areas; each dot represents one mouse; median shown as horizontal grey line inside the box; mean is marked by a grey diamond; unpaired, one-tailed, Welch’s t-test. **(c, d)** MSD immunoassays standardized for tissue lysates; SilencioMA (n=14), aCSF (n = 7; one animal was excluded as an outlier, see Methods). Box plots of the fold difference of total tau and pT231-tau (**c**), and of indicated Aβ peptides (**d**), in cortices from SilencioMA compared to aCSF injected 5XFAD/PS19 mice; unpaired, one-tailed, Mann-Whitney for total tau, Aβ38, Aβ42; unpaired, one-tailed, Welch’s t-test for pT231-tau and Aβ40.

To examine the effect of SilencioMA on tau protein and Aβ peptides we employed quantitative immunoassays based on the Meso Scale Discovery (MSD) electrochemiluminescence platform. We established experimental parameters and dilutions to adjust MSD assays for mouse brain homogenates and ran lysates from posterior cortices of SilencioMA-treated (n=14) and vehicle-only (aCSF)-treated mice (n=7; one outlier excluded, see Methods) (Figure 3c, d, Supplementary Figure 2, Supplementary Table 3). We found that SilencioMA treatment caused a significant reduction of total tau (46.4%, p=0.008) and pT231-tau (59.3%, p=0.046) (Figure 3c), as well as of all three Aβ peptides measured (Aβ38: 35.3%, p=0.019; Aβ40: 55.7%, p=0.033 and Aβ42: 25.5%, p=0.047) (Figure 3d). These findings indicate that SilencioMA can effectively target *MAPT* and *APP* for durable reduction of tau, phospho-tau and Aβ species.

### SilencioMA effectively reduces the development of tau and Aβ pathology *in vivo*

To test the ability of SilencioMA to affect the progression of tau and Aβ neuropathology we stained brain sections of treated and control 5XFAD/PS19 mice with AT8 and NAB228 monoclonal antibodies, 2 months after injection. AT8 is a widely used antibody that recognizes hyperphosphorylated tau (double epitope pSer202/pThr205) and detects NFTs, NTs and other forms of tau pathology [35] [63] [27]. NAB228 recognizes residues 1 to 11 of Aβ peptides and detects amyloid plaques and APP [56] [95]. We found that SilencioMA treated mice showed substantial reduction (60%) in pathological tau throughout the entire brain compared to aCSF controls with the effect approaching statistical significance (p=0.05) (Figure 4a-i; Supplementary Figure 3a-h; Supplementary Table 4). Similarly, we found significant reduction in amyloid plaques in mice treated with SilencioMA versus controls (54%, p=0.04) (Figure 5a-g; Supplementary Figure 3i-l; Supplementary Table 4). Taken together, our data suggest that SilencioMA is efficacious in halting ADNC, making it a unique, dual-targeting compound and a promising candidate for further development as a DMT for AD.

**Figure 4.**
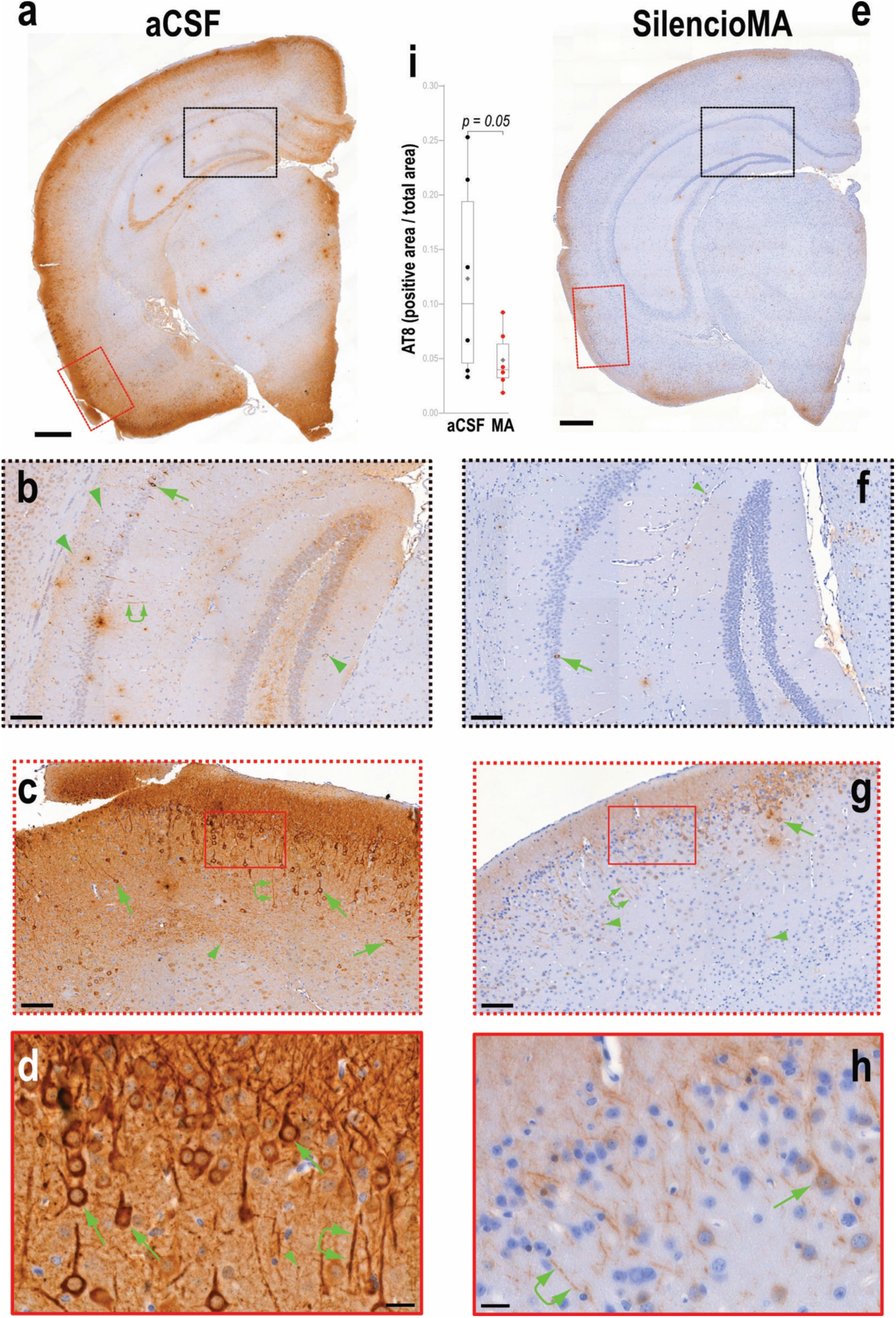
SilencioMA ameliorates tau pathology *in vivo*. Representative coronal hemispheric brain sections of 5XFAD/PS19 mice injected with aCSF (**a-d**) or SilencioMA (**e-h**), immunostained with AT8 (brown chromogen) and counterstained with hematoxylin (blue). Higher magnification images correspond to boxed areas from the hemisphere or image shown above. Numerous NFTs (arrows) and NTs (arrowheads: cross section; linked double arrowheads: longitudinal section) are evident in aCSF (**a-d**), while far fewer are seen in mice treated with SilencioMA (**e-h**); scale bars: a, e =500μm; b, f =50μm; c, g =100μm; d, h =20μm. **(i)** AT8-positive area was quantified from matched coronal hemispheric sections of digitized slides from 5XFAD/PS19 mice treated with SilencioMA (n=6) or aCSF (n=6), normalized to total brain area per hemisphere, and plotted (each dot represents one mouse); unpaired, one-tailed, Welch’s t-test.

**Figure 5.**
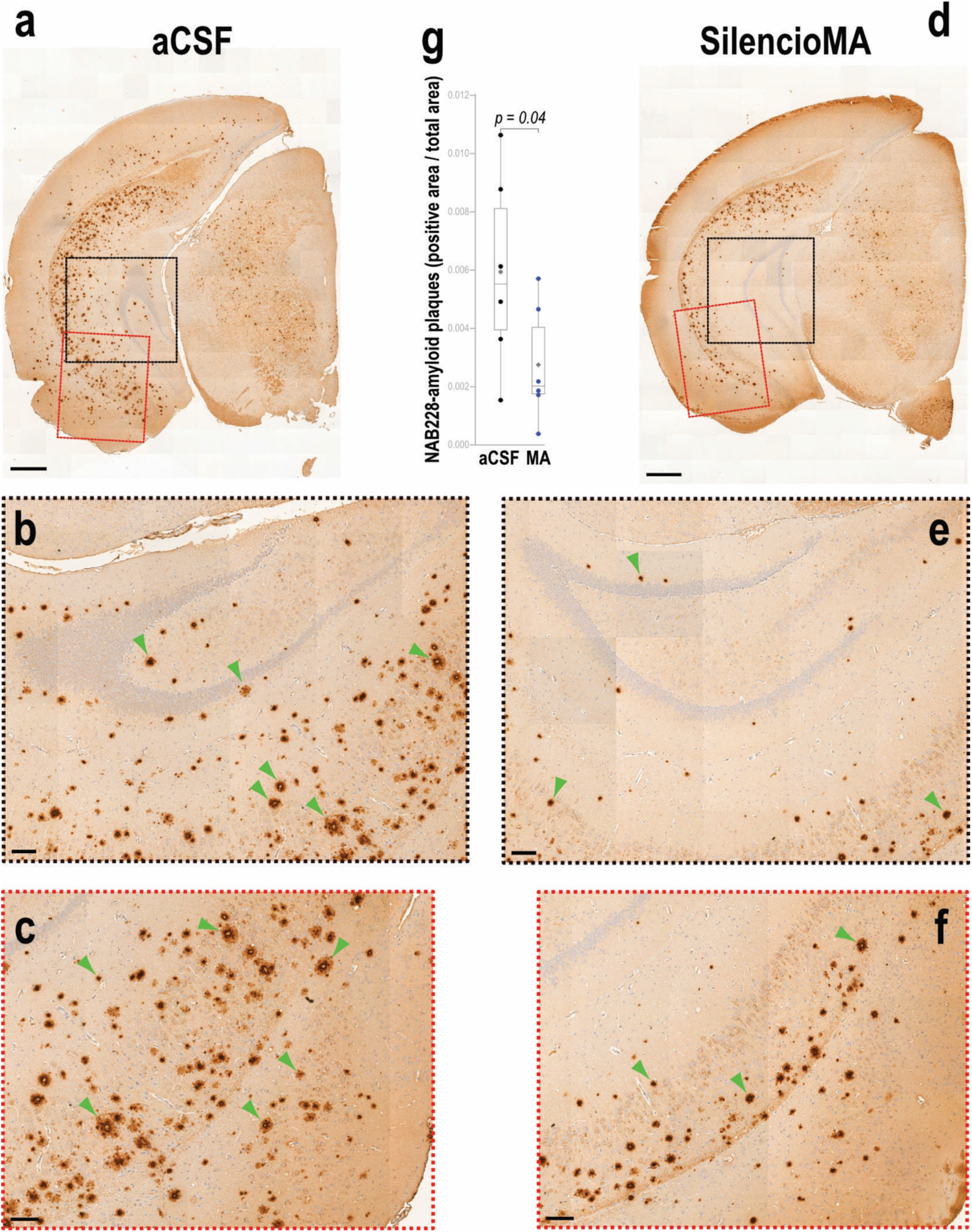
SilencioMA ameliorates amyloid plaque pathology *in vivo*. Representative coronal hemispheric brain sections of 5XFAD/PS19 mice injected with aCSF (**a-c**) or SilencioMA (**d-f**), immunostained with NAB228 (brown chromogen) and counterstained with hematoxylin (blue). Higher magnification images correspond to boxed areas from the hemisphere shown above. Numerous amyloid plaques (arrows) are evident in aCSF (**a-c**), while far fewer are seen in mice treated with SilencioMA (**d-f**); scale bars: a, d = 500μm; b, e =100μm; c, f = 100μm. **(g)** Amyloid plaques were quantified from matched coronal hemispheric sections of digitized slides from 5XFAD/PS19 mice treated with SilencioMA (n=6) or aCSF (n=6), normalized to total brain area per hemisphere, and plotted (each dot represents one mouse); unpaired, one-tailed, Welch’s t-test.

## DISCUSSION

In this study, we describe SilencioMA, a single compound that can engage both major pathological drivers of AD, resulting in significant reductions of *MAPT* and *APP* mRNAs, as well as pathological tau and Aβ species, in a mouse model that develops severe ADNC. A single administration of SilencioMA leads to sustained reduction of ADNC for at least two months post injection. SilencioMA targets conserved regions in human *MAPT* and *APP* mRNA that are found in all isoforms and all major disease variants. The synthesis of SilencioMA is straightforward, its chemical modifications are clinically safe, and delivery does not require specialized formulations. SilencioMA is not expected to trigger ARIA or similar side effects associated with immunotherapies because it inhibits production of pathogenic proteins rather that activate the immune system to clear them.

Limitations of our study, like in most preclinical studies, are introduced by the employment of a transgenic model that relies on overexpression of mutant human transgenes to develop rapid ADNC. Mouse models recapitulate neither the slow progression, over years, nor the full complexity of the human disease. More studies are needed to determine optimal dose and dosing intervals for maximal efficacy and safety and to examine whether the simultaneous knockdown of *MAPT* and *APP* might yield synergistic benefits to allow lower doses, potentially mitigating adverse effects that may manifest with more aggressive, single-target suppression.

A desirable feature of SilencioMA, and of siRNA-based designs in general, is that ligands can be attached to the passenger (sense) strand to achieve targeted delivery, leaving the guide (antisense) strand, which is the pharmacologically active component, untouched [62] [66] [20] [23]. Binding to Transferrin Receptor 1 (TfR1) and the transcytosis it triggers, offers an especially attractive route for carrying therapeutics across the blood brain barrier following systemic administration [46] [28] [32] [83] [90]. A range of TfR1 ligands is currently under development, from antibodies to nucleic acid aptamers, with the most advanced designs having reached clinical trials [4] [90] [73]. Future studies will explore such TfR1 conjugates for systemic administration of SilencioMA.

APOE4 exerts a toxic gain of function in AD and lowering its levels in *APOE4* carriers is a therapeutic goal for AD [92]. In community-based and clinic-based autopsy cohorts, many clinically diagnosed AD patients have at least one co-pathology, such as α-synuclein pathology (Lewy Body Dementia) or TDP-43 pathology (Limbic-predominant age-related TDP-43 encephalopathy) and these co-pathologies often act additively, accelerating cognitive decline and altering the patient’s clinical phenotype [85] [70] [44] [50] [76] [69] [82] [58] [51] [88]. The Silencio approach is well-suited for simultaneous silencing of multiple targets, as the passenger strands are linked in a 5’ to 3’ manner, enabling annealing of guides against independent targets, to generate a single compound. Future studies will focus on further investigating a tri-targeting Silencio that silences *APOE*, in addition to *APP* and *MAPT,* that we have developed (our unpublished data), as well as the development of Silencios that target co-pathologies of AD and other neurodegenerative diseases.

### Conclusions

The pathophysiology of AD is anchored by two defining molecular events: the accumulation of extracellular fibrils of Aβ peptides derived from APP; and the formation of intracellular fibrils of hyperphosphorylated tau (MAPT). These interdependent pathologies prompted us to pursue an approach that addresses more effectively their synergistic nature. We developed SilencioMA as a single compound that simultaneously targets *MAPT* and APP, circumventing limitations of monotherapies. The modular design of Silencios permits conjugation of ligands for transcytosis through the blood brain barrier, allowing for systemic administration. Moreover, this modular feature facilitates development of single compounds for simultaneous silencing of more than two targets, which may prove useful for *APOE4* carriers and for patients with AD, α-synuclein, TDP-43 or other co-pathologies.

## LIST OF ABBREVIATIONS

SilencioMA: <u>Silen</u>cing <u>c</u>onnected interfering <u>o</u>ligonucleotides against *<u>M</u>APT* and *<u>A</u>PP*
MAPT: Microtubule Associated Protein Tau
siRNAs: Small interfering RNAs
AD: Alzheimer’s disease
ADNC: Alzheimer’s Disease Neuropathologic Change
IND: Investigational New Drug
mRNA: Messenger RNA
NFTs: Neurofibrillary tangles
NTs: Neuropil threads
NPs: Neuritic plaques
APP: Amyloid Precursor Protein
APOE: Apolipoprotein E
ASOs: Antisense Oligonucleotides
aCSF: Artificial Cerebrospinal fluid
ICV: Intracerebroventricular
LNPs: Lipid Nanoparticles
NIH: National Institutes of Health
MSD: Meso Scale Discovery
ISH: In situ Hybridization
LNA: Locked Nucleic Acid
IHC: Immunohistochemistry
DIG: Digoxigenin
NBF: Neutral Buffered Formalin
RT: Room temperature
BSA: Bovine Serum Albumin
UPLC: Ultra Performance Liquid Chromatography
MS: Mass Spectrometry
DMT: Disease Modifying Therapeutic
FDA: Food and Drug Administration
IND: Investigational New Drug
SNCA: Synuclein alpha
LATE: Limbic-predominant age-related TDP-43 encephalopathy
LBD: Lewy Body Disease

## ETHICS APPROVAL AND CONSENT TO PARTICIPATE

All procedures in this study were conducted in accordance with the National Institutes of Health (NIH) *Guide for the Care and Use of Laboratory Animals* and were reviewed and approved by the University of Pennsylvania Animal Care and Use Committee.

## CONSENT FOR PUBLICATION

Not applicable. This study does not contain any data from individual human subjects.

## AVAILABILITY OF DATA AND MATERIALS

All data generated or analyzed during this study are included in this published article (and its supplementary information files).

## AUTHORS’ CONTRIBUTIONS

Conceptualization: Z.M., Design and Methodology: Z.M., P. S., N.V. and H.X., Experimentation: P.S., N.V., K.X. and T.B., Data acquisition: P.S. and N.V., Data curation and analysis: P.S. and Z.M., Writing-original draft preparation: P.S., Writing-review and editing: P.S., Z.M., N.V., H.X., V.M-Y.L. and E.B.L., Project administration: P.S. and Z.M., Funding acquisition: Z.M. and E.B.L., Supervision: Z.M.

All authors have read and agreed to the published version of the manuscript.

## ACKNOWLEDGEMENTS

The authors would like to thank Dr. Kurt Brunden for many helpful discussions and comments on the manuscript, the Pathology Core Laboratory (RRID:SCR_028406) at the Children’s Hospital of Philadelphia Research Institute (Daniel Martinez) for providing brain sectioning services, and the Human Immunology Core (HIC, RRID: SCR_022380) at the Perelman School of Medicine at the University of Pennsylvania (Drs Xuming Mao, Honghong Sun, and Eline Luning Prak) for assistance with MSD assays.

---

**Supplementary Figure 1.**
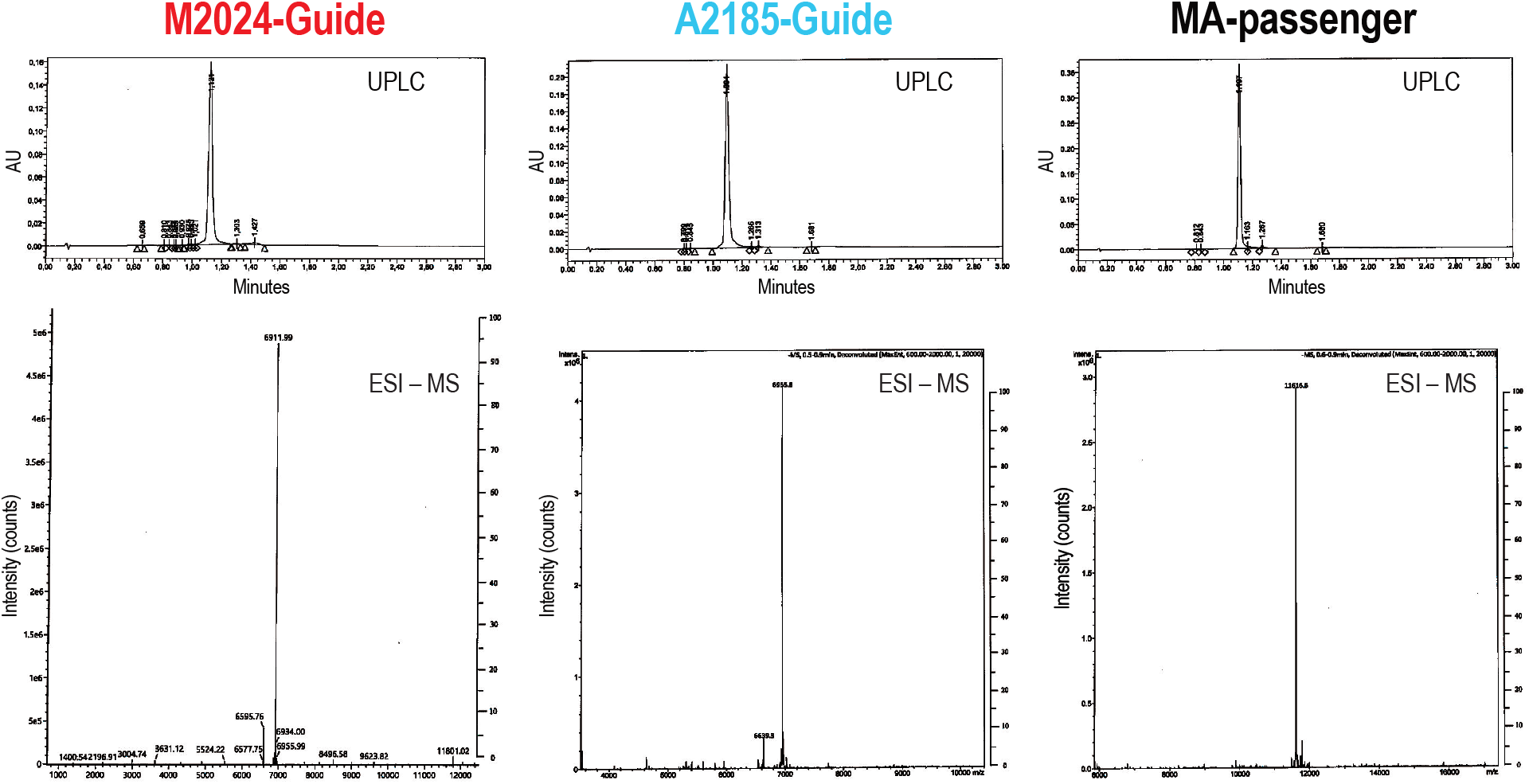
Properties of synthesized oligonucleotides. Ultra Performance Liquid Chromatography (UPLC) indicate the high purity of the oligonucleotides used to generate SilencioMA: M2024-Guide (96.3%), A2185-Guide (97.9%) and MA-passenger (95.2%) Electrospray Ionization Mass Spectrometry (ESI–MS) indicate that the synthesized oligonucleotides are of the expected masses: M2024-Guide (6912.7 Da theoretical, 6911.9 Da observed); A2185-Guide (6955.7 Da theoretical, 6955.8 Da observed); MA-passenger (11616.22 Da theoretical, 11616.6 Da observed).

**Supplementary Figure 2.**
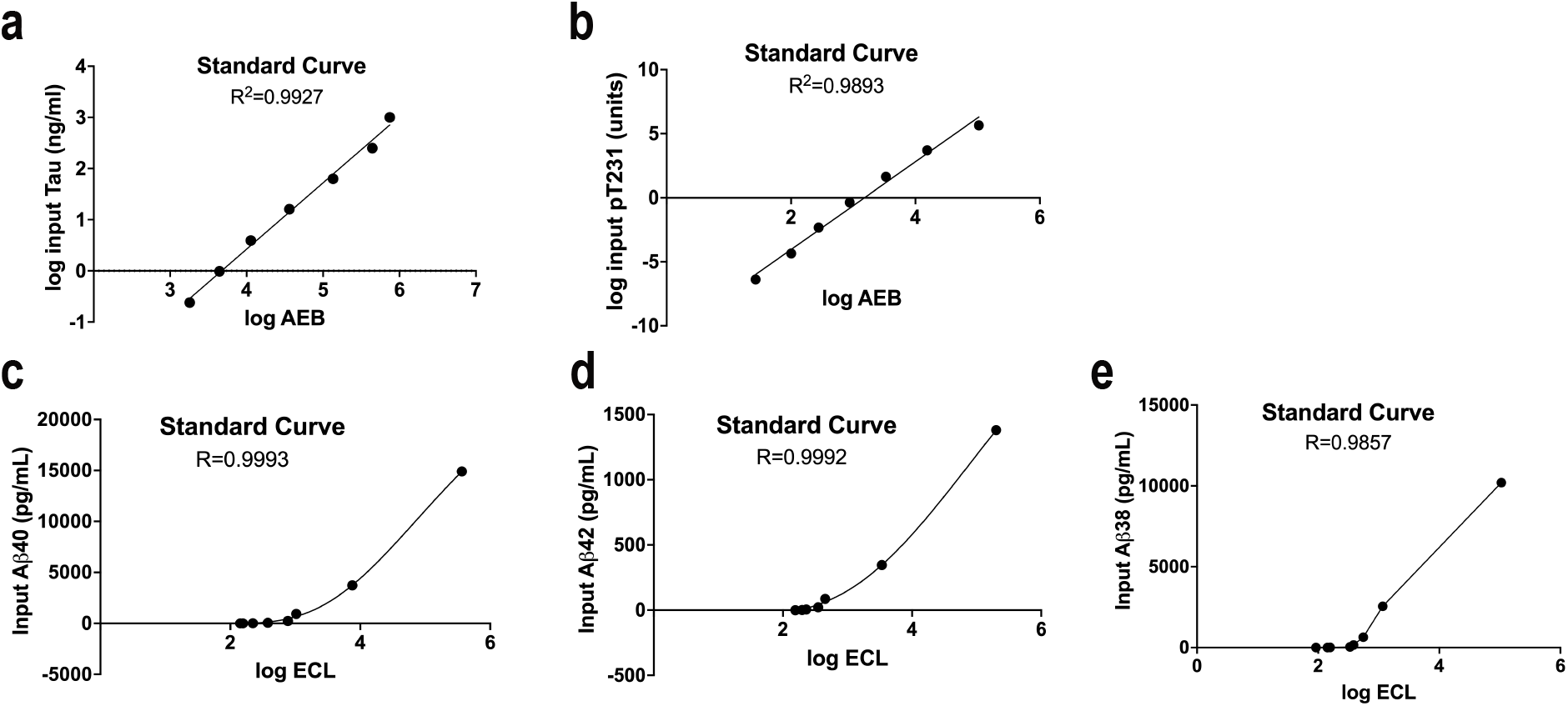
MSD Immunoassays; standard curves. (**a**) Standard curve generated from 7 samples containing known amounts of human tau. (**b**) Standard curve generated from 7 samples containing known amounts of pT231 in relation to total Tau (**c-e**). Standard curves generated from 7 known amounts of Aβ40 (c), Aβ42 (d) and Aβ438 (e).

**Supplementary Figure 3.**
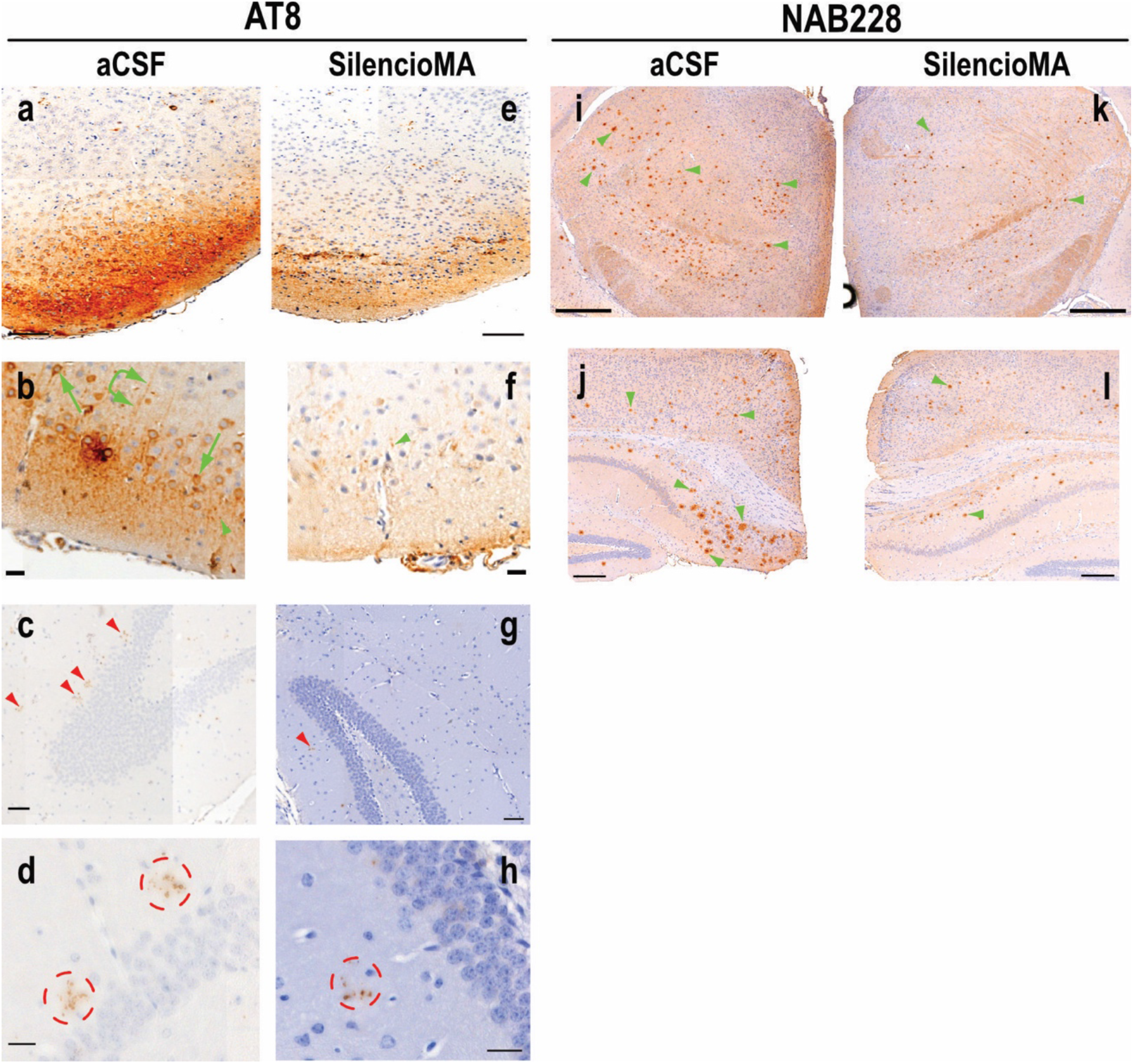
SilencioMA ameliorates ADNPC *in vivo*. Additional examples of AT8 (**a-h**) and NAB228 (**i-l**) staining in various brain areas (cortex, hippocampus, thalamus) of different 5XFAD/PS19 mice treated with aCSF (**a, b, c, d, i, j**) or SilencioMA (**e, f, g, h, k, l**). Neurofibrillary tangles (green arrows) and neuropil threads (green arrowheads: cross section; linked green double arrowheads: longitudinal section) (**a, b**), and neuritic plaques (red arrows or red circles) (**c, d**) are evident in aCSF treated (control) mice while they are reduced in mice treated with SilencioMA (**e - h)**. Amyloid plaques (green arrowheads), evident in aCSF (**i, j**), are reduced in SilencioMA treated mice (**k, l**). Scale bars in a, e = 50 μm; b, f = 20 μm; c, g = 50 μm; d, h= 20 μm; i, k = 500 μm; j, l = 200 μm.

