## Supplementary Table 1 for "SilencioMA: A single compound that targets both *MAPT* and *APP* as a potential disease modifying therapeutic for Alzheimer’s Disease"

| Name | 5′-3′sequence | Target |
|---|---|---|
| MAPT-F | CTTAGCAACGTCCAGTCCAA | MAPT; RT-qPCR in HEK293 cells |
| MAPT-R | TCACCTTGCTCAGGTCAAC | |
| TgMAPT-F | GAAGATTGGGTCCCTGGAC | MAPT transgene transcript; RT-qPCR in 5XFAD/PS19 mice |
| TgMAPT-R | GTCTGTCTTGGCTTTGGCG | |
| HsGAPDH-F | GAGTCCACTGGCGTCTTCAC | GAPDH; RT-qPCR in HEK 293 cells |
| HsGAPDH-R | GAGGCATTGCTGATGATCTTGAG | |
| MmGAPDH-F | CATCACTGCCACCCAGAAGACTG | GAPDH; RT-qPCR in 5XFAD/PS19 mice |
| MmGAPDH-R | ATGCCAGTGAGCTTCCCGTTCAG | |
| APP-F | TTCATCATGGTGTGGTGGAG | APP; RT-qPCR in HEK 293 cells |
| APP-R | CTGCATCTGCTCAAAGAACTTG | |
| TgAPP-F | GTGGAGGTTGACGCCGCT | APP transgene transcript; RT-qPCR in 5XFAD/PS19 mice |
| TgAPP-R | AAAGAACTTGTAGGTTGGATTTTCG | |
| TRW | ACACGTCTCCACGGCATCTCAGC | MAPT for genotyping PS19 mouse |
| MoPrPAs-1 | TCCCCCAGCCTAGACCACGAGAAT | |
| 5xAPPF1 | AGGACTGACCACTCGACCAG | APP for genotyping 5xFAD mouse |
| 5XAPPR1 | CGGGGGTCTAGTTCTGCAT | |
| 5XCTRF1 | CAAATGTTGCTTGTCTGGTG | |
| 5XCTRR1 | GTCAGTCGAGTGCACAGTTT | |
| 5XPS1F1 | AATAGAGAACGGCAGGAGCA | PSEN1 for genotyping 5xFAD mouse |
| 5XPS1R1 | GCCATGAGGGCACTAATCAT | |
| 5XCTRF2 | CTAGGCCACAGAATTGAAAGATCT | |
| 5XCRTR2 | GTAGGTGGAAATTCTAGCATCATCC | |
| RD-F | CATCCCACCTGAGCTCACAGAAAG | RD for genotyping 5xFAD mouse |
| RD-R | GCCTACAACAGAGGAGCTTCTAGC | |
