## Supplementary Table 2 for "SilencioMA: A single compound that targets both *MAPT* and *APP* as a potential disease modifying therapeutic for Alzheimer’s Disease"

| Sample Name | Group | Target Name | Posterior cortex Ct mean | Hippocampus Ct mean |
|---|---|---|---|---|
| 4 | Treated (SilencioMA) | GAPDH | 17.55 | 19.11 |
| 4 | Treated (SilencioMA) | APP | 18.51 | 19.56 |
| 4 | Treated (SilencioMA) | MAPT | 20.58 | 22.99 |
| 11 | Treated (SilencioMA) | GAPDH | 17.94 | 19.48 |
| 11 | Treated (SilencioMA) | APP | 18.64 | 19.94 |
| 11 | Treated (SilencioMA) | MAPT | 19.58 | 21.62 |
| 17 | Treated (SilencioMA) | GAPDH | 17.78 | 19.07 |
| 17 | Treated (SilencioMA) | APP | 18.23 | 18.71 |
| 17 | Treated (SilencioMA) | MAPT | 19.15 | 20.10 |
| 18 | Treated (SilencioMA) | GAPDH | 17.85 | 19.37 |
| 18 | Treated (SilencioMA) | APP | 18.48 | 19.78 |
| 18 | Treated (SilencioMA) | MAPT | 19.38 | 20.86 |
| 27 | Treated (SilencioMA) | GAPDH | 17.86 | 19.16 |
| 27 | Treated (SilencioMA) | APP | 18.45 | 18.71 |
| 27 | Treated (SilencioMA) | MAPT | 19.67 | 20.89 |
| 56 | Treated (SilencioMA) | GAPDH | 17.66 | 18.90 |
| 56 | Treated (SilencioMA) | APP | 20.45 | 23.95 |
| 56 | Treated (SilencioMA) | MAPT | 20.62 | 22.50 |
| 61 | Treated (SilencioMA) | GAPDH | 17.57 | 19.45 |
| 61 | Treated (SilencioMA) | APP | 18.68 | 20.44 |
| 61 | Treated (SilencioMA) | MAPT | 19.70 | 21.52 |
| 66 | Treated (SilencioMA) | GAPDH | 17.29 | 18.94 |
| 66 | Treated (SilencioMA) | APP | 17.91 | 19.45 |
| 66 | Treated (SilencioMA) | MAPT | 18.49 | 20.48 |
| 90 | Treated (SilencioMA) | GAPDH | 17.87 | 19.29 |
| 90 | Treated (SilencioMA) | APP | 18.16 | 19.68 |
| 90 | Treated (SilencioMA) | MAPT | 19.66 | 21.34 |
| 94 | Treated (SilencioMA) | GAPDH | 17.61 | 19.29 |
| 94 | Treated (SilencioMA) | APP | 17.90 | 19.80 |
| 94 | Treated (SilencioMA) | MAPT | 19.48 | 21.82 |
| 95 | Treated (SilencioMA) | GAPDH | 17.56 | 19.26 |
| 95 | Treated (SilencioMA) | APP | 17.56 | 20.27 |
| 95 | Treated (SilencioMA) | MAPT | 19.40 | 21.86 |
| 99 | Treated (SilencioMA) | GAPDH | 17.77 | 19.33 |
| 99 | Treated (SilencioMA) | APP | 17.81 | 19.70 |
| 99 | Treated (SilencioMA) | MAPT | 19.03 | 21.20 |
| 118 | Treated (SilencioMA) | GAPDH | 17.65 | 19.85 |
| 118 | Treated (SilencioMA) | APP | 18.23 | 21.35 |
| 118 | Treated (SilencioMA) | MAPT | 18.94 | 21.95 |
| 119 | Treated (SilencioMA) | GAPDH | 17.85 | 19.37 |
| 119 | Treated (SilencioMA) | APP | 18.22 | 20.05 |
| 119 | Treated (SilencioMA) | MAPT | 19.75 | 21.47 |
| 148 | Control (aCSF) | GAPDH | 17.40 | 19.14 |
| 148 | Control (aCSF) | APP | 17.71 | 19.27 |
| 148 | Control (aCSF) | MAPT | 18.49 | 20.65 |
| 153 | Control (aCSF) | GAPDH | 18.37 | 18.93 |
| 153 | Control (aCSF) | APP | 18.39 | 18.59 |
| 153 | Control (aCSF) | MAPT | 19.84 | 20.95 |
| 158 | Control (aCSF) | GAPDH | 18.16 | 19.22 |
| 158 | Control (aCSF) | APP | 17.92 | 19.55 |
| 158 | Control (aCSF) | MAPT | 19.90 | 21.07 |
| 163 | Control (aCSF) | GAPDH | 18.22 | 18.76 |
| 163 | Control (aCSF) | APP | 18.14 | 18.99 |
| 163 | Control (aCSF) | MAPT | 19.32 | 20.51 |
| 165 | Control (aCSF) | GAPDH | 17.80 | 19.04 |
| 165 | Control (aCSF) | APP | 18.14 | 18.83 |
| 165 | Control (aCSF) | MAPT | 19.07 | 20.48 |
| 171 | Control (aCSF) | GAPDH | 17.80 | 18.80 |
| 171 | Control (aCSF) | APP | 17.72 | 18.81 |
| 171 | Control (aCSF) | MAPT | 19.21 | 20.76 |
| 181 | Control (aCSF) | GAPDH | 17.47 | 18.28 |
| 181 | Control (aCSF) | APP | 18.15 | 18.37 |
| 181 | Control (aCSF) | MAPT | 18.69 | 19.57 |
| 200 | Control (aCSF) | GAPDH | 17.34 | 18.79 |
| 200 | Control (aCSF) | APP | 17.73 | 19.04 |
| 200 | Control (aCSF) | MAPT | 18.58 | 20.58 |
