## Supplementary Table 3 for "SilencioMA: A single compound that targets both *MAPT* and *APP* as a potential disease modifying therapeutic for Alzheimer’s Disease"

| Treatment | pT231 (pg/mL) of Tau - mean | Total Tau (ng/mL) - mean | Absolute pT231 (pg/mL) - mean |
|---|---|---|---|
| SilencioMA | 255.8830725 | 7.783095688 | 1991.562438 |
| SilencioMA | 855.6163729 | 49.43329741 | 42295.93863 |
| SilencioMA | 1284.96282 | 65.57018901 | 84255.25497 |
| SilencioMA | 659.5374583 | 36.59222431 | 24133.94261 |
| SilencioMA | 573.3197428 | 40.41596112 | 23171.26844 |
| SilencioMA | 283.0137628 | 9.154013177 | 2590.711714 |
| SilencioMA | 1237.690907 | 51.40554446 | 63624.17496 |
| SilencioMA | 1548.461405 | 61.90851548 | 95862.94686 |
| SilencioMA | 1459.413747 | 62.90604846 | 91805.95191 |
| SilencioMA | 1070.259718 | 57.94139868 | 62012.345 |
| SilencioMA | 1008.430862 | 63.74241297 | 64279.81648 |
| SilencioMA | 1631.649285 | 62.65109582 | 102224.6157 |
| SilencioMA | 783.8434474 | 28.85038964 | 22614.18887 |
| SilencioMA | 687.738106 | 47.76083172 | 32846.94394 |
| aCSF | 2249.539255 | 126.8130065 | 285270.8363 |
| aCSF | 916.2152515 | 51.87860271 | 47531.96703 |
| aCSF | 734.816281 | 31.33245411 | 23023.5974 |
| aCSF | 2148.443739 | 105.8619333 | 227438.4078 |
| aCSF | 831.7227566 | 83.78795809 | 69688.35147 |
| aCSF | 1307.282227 | 92.56761575 | 121011.9989 |
| aCSF | 3696.559795 | 366.3622482 | 1354279.957 |
| aCSF | 925.082358 | 110.7983127 | 102497.5644 |

| Sample # | Abeta38 (pg/mL) | | | Abeta40 (pg/mL) | | | Abeta42 (pg/mL) | | |
|---|---|---|---|---|---|---|---|---|---|
| Treatment | Dup 1 | Dup 2 | Mean | Dup 1 | Dup 2 | Mean | Dup 1 | Dup 2 | Mean |
| **SilencioMA** | 201.0707 | 225.0235 | **213.0471** | 45.0591 | 37.2093 | **41.1342** | 390.8061 | 386.2627 | **388.5344** |
| **SilencioMA** | 370.5176 | 501.1579 | **435.8378** | 2057.2524 | 2086.2353 | **2071.7439** | 1027.9988 | 1069.0351 | **1048.5170** |
| **SilencioMA** | 391.4957 | 645.9196 | **518.7076** | 1659.8608 | 1753.3943 | **1706.6276** | 1013.2932 | 1106.3995 | **1059.8463** |
| **SilencioMA** | 381.0495 | 248.5271 | **314.7883** | 984.8636 | 1156.5645 | **1070.7141** | 989.4234 | 1056.0142 | **1022.7188** |
| **SilencioMA** | 784.3757 | 792.1003 | **788.2380** | 2194.0058 | 2310.8531 | **2252.4294** | 1261.6625 | 1247.5385 | **1254.6005** |
| **SilencioMA** | 201.0707 | 213.1045 | **207.0876** | 37.2093 | 29.5132 | **33.3613** | 62.8019 | 60.8305 | **61.8162** |
| **SilencioMA** | 327.5010 | 510.6845 | **419.0927** | 460.3009 | 424.9079 | **442.6044** | 699.1562 | 669.9708 | **684.5635** |
| **SilencioMA** | 338.3918 | 422.3350 | **380.3634** | 745.3383 | 772.3861 | **758.8622** | 800.5953 | 807.3990 | **803.9971** |
| **SilencioMA** | 520.1424 | 510.6845 | **515.4134** | 1644.6511 | 1538.7375 | **1591.6943** | 873.5641 | 840.3604 | **856.9623** |
| **SilencioMA** | 815.0188 | 776.6076 | **795.8132** | 3137.3263 | 2463.1345 | **2800.2304** | 1190.0397 | 1128.4422 | **1159.2409** |
| **SilencioMA** | 520.1424 | 481.8945 | **501.0185** | 1822.6045 | 1891.7824 | **1857.1935** | 1051.9285 | 1063.4491 | **1057.6888** |
| **SilencioMA** | 442.4931 | 510.6845 | **476.5888** | 1945.5980 | 1911.5984 | **1928.5982** | 1082.1948 | 1034.1581 | **1058.1765** |
| **SilencioMA** | 349.1905 | 338.3918 | **343.7911** | 441.6253 | 537.5867 | **489.6060** | 728.1097 | 781.6596 | **754.8846** |
| **SilencioMA** | 327.5010 | 422.3350 | **374.9180** | 1367.8425 | 1495.0964 | **1431.4694** | 1014.4981 | 1042.0553 | **1028.2767** |
| **aCSF** | 391.4957 | 452.4559 | **421.9758** | 665.7879 | 695.7433 | **680.7656** | 753.5133 | 782.9818 | **768.2476** |
| **aCSF** | 837.5652 | 784.3757 | **810.9704** | 3778.9986 | 3944.4405 | **3861.7196** | 1336.5296 | 1367.8871 | **1352.2083** |
| **aCSF** | 815.0188 | 852.3959 | **833.7074** | 4323.4996 | 4267.0636 | **4295.2816** | 1416.8932 | 1395.2169 | **1406.0551** |
| **aCSF** | 874.3512 | 895.9683 | **885.1598** | 5135.1780 | 5227.0432 | **5181.1106** | 1423.9826 | 1459.3386 | **1441.6606** |
| **aCSF** | 271.5992 | 349.1905 | **310.3948** | 1295.6385 | 1169.8725 | **1232.7555** | 903.9988 | 882.7077 | **893.3533** |
| **aCSF** | 799.7820 | 917.2587 | **858.5204** | 4642.1933 | 4630.9319 | **4636.5626** | 1420.3922 | 1447.1365 | **1433.7644** |
| **aCSF** | 529.5328 | 679.7877 | **604.6603** | 2390.3436 | 2406.2535 | **2398.2985** | 1107.1892 | 1131.2992 | **1119.2442** |
| **aCSF** | 815.0188 | 662.9598 | **738.9893** | 1002.5409 | 894.0929 | **948.3169** | 912.3268 | 915.9271 | **914.1270** |
