## Supplementary Table 4 for "SilencioMA: A single compound that targets both *MAPT* and *APP* as a potential disease modifying therapeutic for Alzheimer’s Disease"

| Total area (µm^2) | AT8: Positive area (µm^2) | Treatment |
|---|---|---|
| 72620970.5 | 4848314 | aCSF |
| 41436498.6 | 1371977.75 | aCSF |
| 33059328.3 | 7075863.5 | aCSF |
| 24944069.4 | 6311763.5 | aCSF |
| 29358961.1 | 1143889.625 | aCSF |
| 63363206.5 | 8478303 | aCSF |
| 74069400.1 | 5217826.5 | SilencioMA |
| 33592364.3 | 1029880.75 | SilencioMA |
| 39452352.5 | 3642594.8 | SilencioMA |
| 24815368.3 | 1047436.75 | SilencioMA |
| 25017745 | 467298.1875 | SilencioMA |
| 81664960.5 | 3054269.5 | SilencioMA |

| Total area (μm^2) | NAB228: Positive area (μm^2) | Treatment |
|---|---|---|
| 35308058.4 | 173550.7 | aCSF |
| 33433658.7 | 355574 | aCSF |
| 29509195.4 | 45374.2539 | aCSF |
| 22662709.1 | 82324.7 | aCSF |
| 26664696 | 234016.8438 | aCSF |
| 39013618.3 | 239021.7 | aCSF |
| 32411639.6 | 184998.1 | SilencioMA |
| 28829970.3 | 134275.0781 | SilencioMA |
| 25388374.5 | 47255.4414 | SilencioMA |
| 29343730.8 | 50285.3 | SilencioMA |
| 21000686.1 | 45752.9219 | SilencioMA |
| 32766878.8 | 12475.4756 | SilencioMA |
